# IGF1R undergoes active and directed centripetal transport on filopodia upon receptor activation

**DOI:** 10.1101/754960

**Authors:** Denis Krndija, Michael Fairhead

## Abstract

Filopodia are thin, actin-based membrane protrusions with roles in sensing external mechanical and chemical cues, such as growth factor gradients in tissues. It was proposed that the chemical sensing role of filopodia is achieved through clearance of activated signaling receptors from filopodia. Type I insulin-like growth factor receptor (IGF1R) is a key regulator of normal development and growth, as well as tumor development and progression. Its biological roles depend on its activation upon IGF1 binding at the cell membrane. IGF1R behavior at the cell membrane and in particular in filopodia, has not been established. We found that IGF1 activation led to a gradual reduction in IGF1R puncta in filopodia, and that this clearance depended on actin, non-muscle myosin II, and IGF1R kinase activity. Using single particle tracking of filopodial IGF1R, we established that ligand-free IGF1R undergoes non-directional unidimensional diffusion along the filopodium. Moreover, after initial diffusion, the ligand-bound IGF1R is actively transported along the filopodium towards the filopodium base, and consequently cleared from the filopodium. Our results show that IGF1R can move directionally on the plasma membrane protrusions, supporting a sensory role for filopodia in interpreting local IGF1 gradients.

## Introduction

Filopodia are thin membrane protrusions of 100 to 200 nm diameter, containing 10-30 actin filaments bundled together by actin cross-linking proteins (Ridley, 2011; Zhuravlev et al., 2010). The structure of filopodia is established dynamically: F-actin in filopodia undergoes retrograde flow, as G-actin monomers polymerize at the tip of filopodia and non-muscle myosin II moves F-actin bundles towards the cell body (Mattila and Lappalainen, 2008). The filopodium can maintain a stable shape amidst this flux through integrin attachment at the tip (Zhang et al., 2004), leading to establishment of a focal adhesion (Partridge and Marcantonio, 2006). Filopodia have roles in sensing external mechanical and chemical cues, in cell migration and cell-cell interactions (Mattila and Lappalainen, 2008), and are important for processes such as blood vessel formation (Lawson and Weinstein, 2002), neuron pathfinding (Dent and Gertler, 2003), wound healing (Noselli, 2002), and cancer metastasis (Machesky, 2008). The role of filopodia in sensing chemical cues has been proposed in various systems, including angiogenic sprouting in mouse retinal development (Gerhardt et al., 2003), tracheal and wing disc morphogenesis in Drosophila (Ribeiro et al., 2002; Roy et al., 2011) and primordial germ cell polarization and migration in zebrafish (Meyen et al., 2015). Clearance of activated signaling receptors from filopodia has been suggested as a potential mechanism for the proposed sensing role of filopodia (Lidke et al., 2005).

The insulin-like growth factor receptor (IGF1R) plays diverse roles in development, growth, cell survival, and tumor invasion (Chitnis et al., 2008; LeRoith and Roberts, 2003). IGF1R is a tyrosine kinase membrane receptor and a therapeutic target for antibodies and small molecule kinase inhibitors (Yuen and Macaulay, 2008). IGF1R is activated upon binding of circulating IGF1 (or IGF2), which is predominantly released by the liver but also by other tissues, including muscle and cartilage (Laron, 2001). Upon ligand-dependent autophosphorylation, IGF1R initiates downstream signaling, particularly through the phosphoinositide 3-kinase (PI3K) and mitogen-activated protein (MAP) kinase pathways (Chitnis et al., 2008). Clearance of the activated receptor from the plasma membrane occurs through IGF1R internalization via clathrin-coated pits and caveolae (Matthews et al., 2008; Sehat et al., 2008).

IGF1 can act both as an endocrine hormone and as a paracrine hormone (Laron, 2001), or even in an autocrine manner when secreted by cancer cells (Yuen and Macaulay, 2008). IGF1 levels are not homogeneous around cells in the tissue—gradients of more than 10-fold difference in IGF concentration have been measured in human tissues (Desvigne et al., 2005). Apart from the IGF1 secretion itself, gradients of IGF binding proteins (IGFBP) (Holly and Perks, 2006), as found in the placenta (Han et al., 1996), could also regulate the spatial complexity of IGF1R signaling. The importance of IGF1 in the development of the central nervous system, especially in neuronal migration and positioning (Hurtado-Chong et al., 2009), and in post-wounding axonal sprouting (Guthrie et al., 1995) could be associated to IGF1 spatial gradients. These gradients could elicit cellular responses—IGF1 signaling has been shown to directly promote cell migration by stimulating membrane protrusions and directed cell migration (Guvakova et al., 2002; Haase et al., 2003). The IGF1 receptor dynamics at the plasma membrane remain, however, poorly understood. In particular, its role and behavior in filopodia has never been addressed before. Here we investigated how IGF1R distribution in filopodia changed in response to receptor activation, how these changes were mediated, and then characterized the dynamics of ligand-free and ligand-bound IGF1R using single particle tracking.

## Results and Discussion

### IGF1 induces the clearance of IGF1R from filopodia in an actin- and non-muscle myosin II-dependent manner

We studied the IGF1R distribution in filopodia in a human prostate carcinoma cell line (DU145) (Stone et al., 1978)—these cells express IGF1R and respond to IGF1 by increasing their invasive capacity (Saikali et al., 2008). IGF1R staining in DU145 filopodia exhibited a punctate distribution (**Fig. 1A**), and treatment with IGF1 for 30 minutes decreased the number of IGF1R puncta in filopodia (**Fig. 1A**). To further characterize this process, IGF1R signal in filopodia was measured at various times after IGF1 treatment (**Fig. 1B**). Filopodial IGF1R signal declined gradually, with ∼37% depletion at 10 min and ∼53% depletion at 30 min (Fig. 1B). To establish whether IGF1R tyrosine kinase activity is required for receptor clearance from filopodia, we treated cells with AZ12253801 (AZ), a small molecule inhibitor of IGF1R (Aleksic et al., 2010). AZ fully inhibited the IGF1-induced filopodial clearance of IGF1R (**Fig. 1C**), suggesting IGF1 clearance in the filopodia requires IGF1R signaling. Receptor trafficking in specialized membrane protrusions such as filopodia can be achieved by tethering to actin cytoskeleton and myosin motors, as shown for integrins and myosin X (Zhang et al., 2004). We therefore hypothesized that IGF1R filopodial clearance could be driven by F-actin, which makes up the core of the filopodium. To explore the role of actin in the movement of IGF1R, we treated cells with cytochalasin D, which inhibits F-actin polymerization. Cytochalasin D blocked IGF-induced clearance (**Fig. 1D, Fig. S1A**), indicating that actin polymerization is required for IGF1R transport in filopodia. The continuous transport of actin networks from the leading edge towards the cell body—actin retrograde flow—is maintained by non-muscle myosin II motor anchored near the ‘minus’ end, pulling the filaments inward (Medeiros et al., 2006; Mitchison and Kirschner, 1988; Ponti et al., 2004). Myosin II was shown to localize to the base of filopodia and to be required for the retrograde transport of viruses on filopodia (Lehmann et al., 2005), implying a role for myosin II-mediated contractility in retrograde actin flow in filopodia. To test the involvement of myosin II in IGF1R dynamics, we inhibited myosin II with blebbistatin, and this completely blocked IGF1R clearance (**Fig. 1E, Fig. S1B**). Receptor clearance in filopodia seems thus to depend on both actin polymerization and myosin II contractility. As a control whether any cytoskeletal change was disruptive, we inhibited microtubule polymerization by using nocodazole. Nocodazole treatment had no effect on IGF1R clearance (**Fig. 1F, Fig. S1C**), consistent with the absence of microtubules from filopodia (Machesky, 2008; Mattila and Lappalainen, 2008) and indicating that IGF1R clearance was not sensitive to changes in microtubule stability in the cell. Filopodia are too narrow for endocytosis and do not contain clathrin (Mattila and Lappalainen, 2008), but to determine whether the clearance of IGFR from filopodia depends on endocytosis from other domains of the plasma membrane, cells were treated with dynasore, a dynamin inhibitor that prevents clathrin-mediated and caveolar endocytosis (Macia et al., 2006). IGF1 treatment still induced efficient clearance of IGF1R from filopodia in the presence of dynasore (**Fig. 1G**), suggesting that IGF1R clearance does not depend on endocytosis. Importantly, inhibitor treatments did not alter the levels of inactive IGF1R in filopodia, which were similar to the values obtained in control cells (**Fig. 1C-G**).

**Fig. 1.**
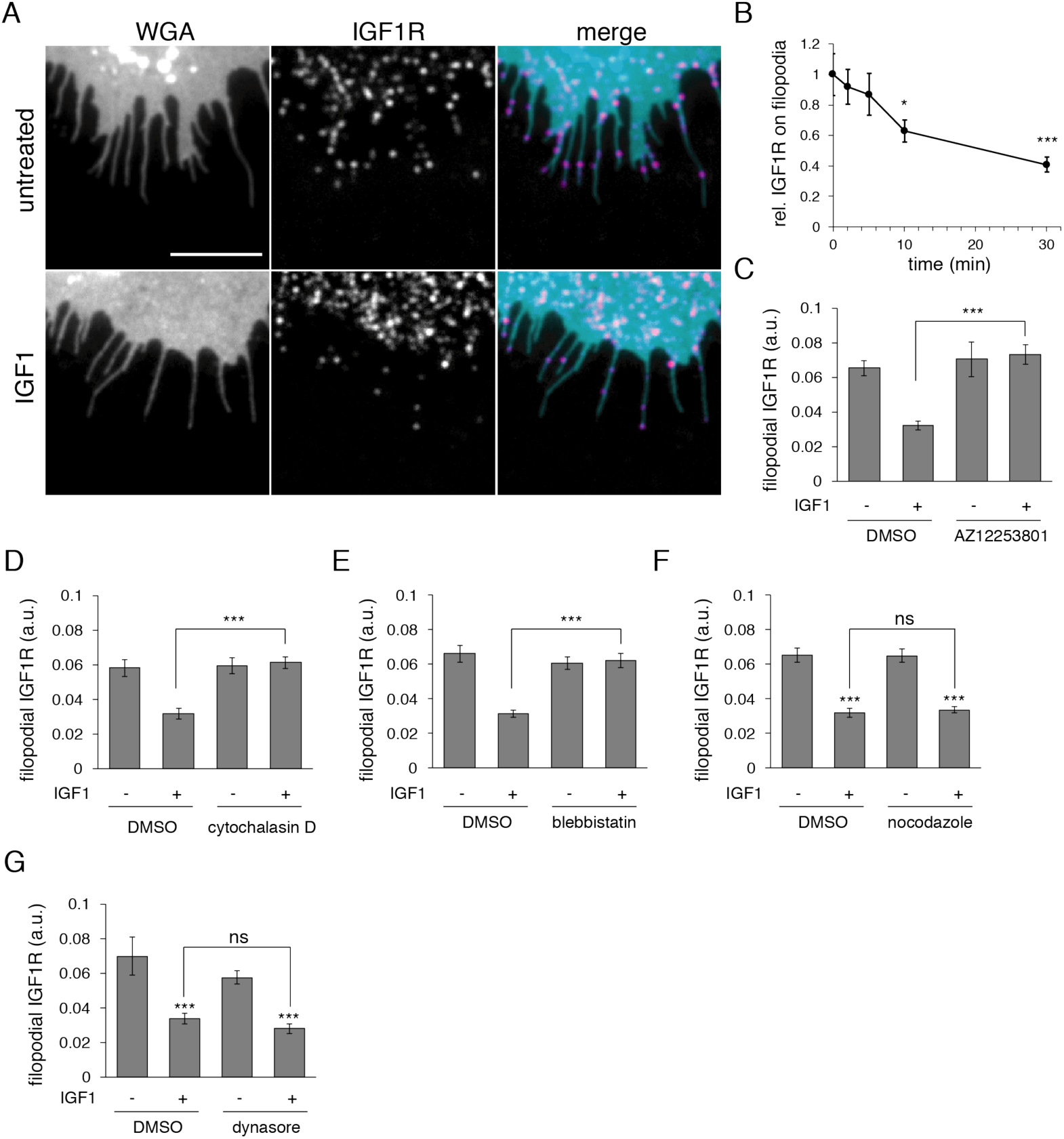
IGF1-induced clearance of IGF1R from filopodia requires actin polymerization and myosin II activity. (A) DU145 cells were serum-starved overnight. Cells in the top panel were untreated and in the bottom panel treated with IGF1 for 30 min. Wide-field fluorescence images are shown of filopodia stained with wheat germ agglutinin (WGA) (left in grayscale) and anti-IGF1R antibody (middle in grayscale), with the merge shown on the right (WGA in cyan, IGF1R in magenta). Scale bar, 5 µm. (B) Filopodial IGF1R staining quantification, untreated or at varying times after adding IGF1 (mean ± s.e.m, N ≥ 28, 2 independent experiments; p<0.0001 for one-way ANOVA followed by Tukey’s test; *** p<0.001, * p<0.05). (C) DU145 cells were serum-starved overnight and treated with IGF1R inhibitor AZ12253801 or DMSO (vehicle control), before adding IGF1 for 30 min and staining for IGF1R intensity on filopodia. AZ12253801 abolished the IGF1-mediated decrease in filopodial IGF1R signal (mean ± s.e.m.; N ≥ 98; 2 independent experiments. p<0.0001 for one-way ANOVA followed by Tukey’s test; *** P<0.001). (D) Cytochalasin D abolished the IGF1-mediated decrease in filopodial IGF1R (mean ± s.e.m.; N ≥ 131; 2 independent experiments. p<0.0001 for one-way ANOVA followed by Tukey’s test; *** P<0.001). (E) Blebbistatin eliminated the IGF1-mediated decrease in filopodial IGF1R signal (mean ± s.e.m.; N ≥ 173; 2 independent experiments. p<0.0001 for one-way ANOVA followed by Tukey’s test; *** P<0.001). (F) Nocodazole did not stop IGF1-induced clearance of filopodial IGF1R (mean ± s.e.m.; N ≥ 155; ns, non-significant; 2 independent experiments. p<0.0001 for one-way ANOVA followed by Tukey’s test). (G) Inhibition of endocytosis did not abrogate IGF1-mediated decrease in filopodial IGF1R signal. Cells were pre-treated with dynasore or with DMSO vehicle control. IGF1 was added for 30 min or left untreated. Cell-surface IGF1R was stained as above (mean ± s.e.m.; N ≥ 67; 2 independent experiments. p<0.0001 for one-way ANOVA followed by Tukey’s test; *** P<0.001; ns, non-significant).

### Ligand-free IGF1R undergoes diffusive motion on filopodia

IGF1R detection using immunofluorescence implies averaging behavior of many molecules, and real-time imaging of the IGF1R movement on filopodia has never been performed. To understand the dynamics of inactive, ligand-free IGF1R in filopodia, we employed live cell imaging and single particle tracking, using a specific affibody (a monomeric non-immunoglobulin binding domain) against IGF1R (Affi_IGF1R_). This affibody binds IGF1R with high affinity and does not activate the receptor (Li et al., 2010). Affi_IGF1R_ was coupled with an acceptor peptide (AP) to enable site-specific biotinylation (bio-Affi_IGF1R_) (Fairhead et al., 2014) and subsequent labelling. We validated the specific binding of bio-Affi_IGF1R_ to the receptor using HeLa cells transfected with IGF1R (**Fig. S2A**). To image ligand-free IGF1R, DU145 cells were labelled with bio-Affi_IGF1R_ followed by streptavidin-labelled quantum dots (QDs) (**Fig. 2A**). The labelling was performed at low concentrations, in order to have one QD-bio-Affi complex per filopodium and facilitate single particle tracking. Three representative receptor trajectories from three independent experiments were overlaid on the corresponding QD-bio-Affi_IGF1R_ images (projections) (**Fig. 2B**). Displacement of the representative tracks shown in Figure 2B, plotted against time (Y-time plot), reveals that there is no systematic directional movement of the receptor during the observation period (**Fig. 2C**). The nature of the ligand-free IGF1R motion was further analyzed by plotting mean squared displacement (MSD) versus time, which facilitates the classification of particle motion into free diffusion (straight-line for MSD v. time), active transport (MSD v. time curves up) and restricted diffusion (MSD v. time curves down) (Saxton and Jacobson, 1997). MSD versus time plot of these trajectories gave an initial straight line, which later levelled off, consistent with confined diffusion (**Fig. 2D**). The initial rate of diffusion, fit to a straight line to give a diffusion coefficient, showed heterogeneity in the diffusion rate of individual receptors (**Fig. 2D**). The mean receptor diffusion rate (0.032 ± 0.006 µm^2^s^-1^, N = 9) is close to the reported diffusion rates for EGFR dimer (Chung et al., 2010). Analysis of directionality confirmed that the movement of IGF1R on filopodia is random, i.e. with a very low directionality index (0.018 ± 0.004, N = 9).

**Fig. 2.**
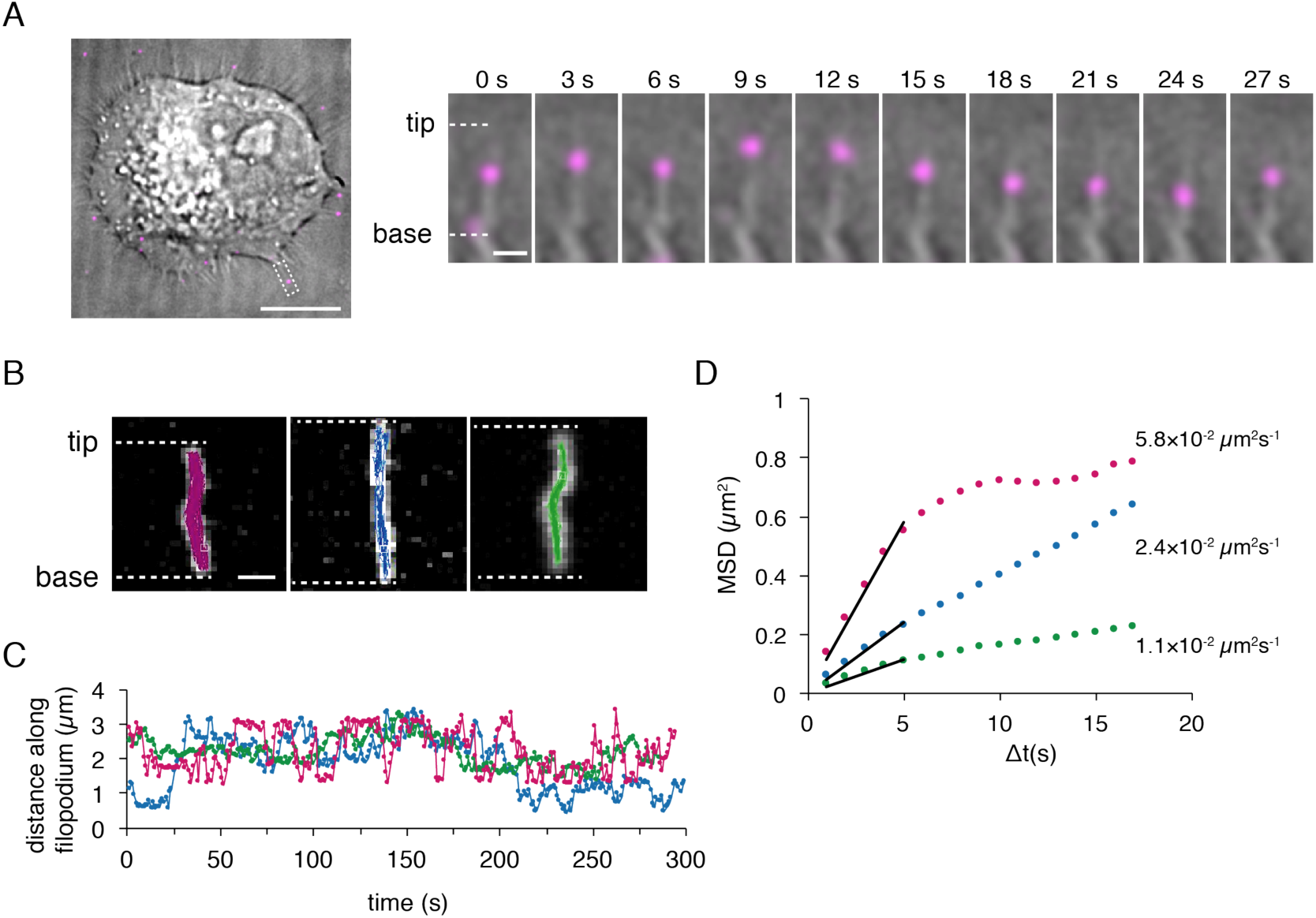
Ligand-free IGF1R displayed diffusive behavior on filopodia. IGF1R on DU145 cells was labelled with biotinylated Affi_IGF1R_, followed by streptavidin-QDs. (A) *Left*: an overlay of Affi-QD (magenta) with the brightfield image (grayscale)—a representative image. Scale bar, 10 µm. *Right*: Ligand-free IGF1R motion on a filopodium (from the boxed region in the left image) at successive indicated times. Scale bar, 2 µm. (B) Full-length Affi-QD trajectories from 3 independent experiments (purple, blue and green) are shown overlaid onto the Affi-QD image stack (grayscale). Scale bar, 2 µm. (C) Y-time plots of the corresponding trajectories showing movement along filopodial axis during the first 300 s of imaging. (D) MSD analysis of Affi_IGF1R_-QD motion on filopodia. MSD values for the tracks shown in (B) were plotted. The first 5 points of the MSD plots were fit linearly to MSD=2DΔt (black lines) to calculate initial diffusion rates, indicated on the graph.

### Ligand-bound IGF1R undergoes directed centripetal transport on filopodia

To understand the behavior of IGF1R on filopodia upon IGF1 binding, the motion of activated IGF1R was analyzed using biotinylated IGF1 (bio-IGF1). It was first determined whether bio-IGF1 bound specifically to IGF1R and not to other potential IGF1 receptors such as integrins (Fujita et al., 2012), using the Affi_IGF1R_ affibody (**Fig. S2B**). After pre-treating DU145 cells with a non-biotinylated Affi_IGF1R_ (to block IGF1R), bio-IGF1 binding was abolished, suggesting that bio-IGF1 can be used to specifically label IGF1R (**Fig. S2C**). For receptor tracking in real time, cell surface-bound bio-IGF1 was detected using streptavidin-QDs, both at low concentrations, to minimize potential receptor/QD clustering. Imaging and tracking of single QDs bound to IGF1 and activated IGF1R on filopodia revealed a directional centripetal movement of IGF1R towards the cell body, resulting in its clearance from the filopodium (**Fig. 3A**). The directional movement was preceded by a period of random movement, as shown by three sample traces, with the trajectories overlaid on the corresponding image projections (**Fig. 3B**) plotted against time (Y-time plot; **Fig. 3C**). The onset of centripetal directional movement was determined from time plots as a continuous directional movement, and was marked with arrows in Figure 4C. Interestingly, the fluctuating speeds of the representative IGF1-QDs undergo a transition to lower and more uniform speed values upon the onset of the centripetal movement, with occasional pausing (**Fig. 3D**). IGF1-QD speed during the directional centripetal movement ranged 0-45 nm/s, with average speed of 12.2 ± 0.003 nm/s. MSD versus time curve analysis of the ligand-bound IGF1R motion (**Fig. 3E**) revealed a clear distinction between a period of random motion showing free diffusion, followed by a period of centripetal movement, which can be characterized as active transport. To confirm the nature of this active transport, we analyzed the directionality of motion, i.e. whether the ligand-bound receptor was as likely to move towards (centripetal motion) or away from the cell body. We found a 9-fold difference in directionality index between these two phases of activated IGF1R transport, showing a clear directionality of centripetal movement (**Fig. 3F**). Taken together, our results underline that the directional centripetal movement of IGF1R on filopodia is dependent on receptor activation upon IGF1 binding; furthermore, we uncovered that this is an active transport, in contrast to the diffusive behavior of the ligand-free receptor.

**Fig. 3.**
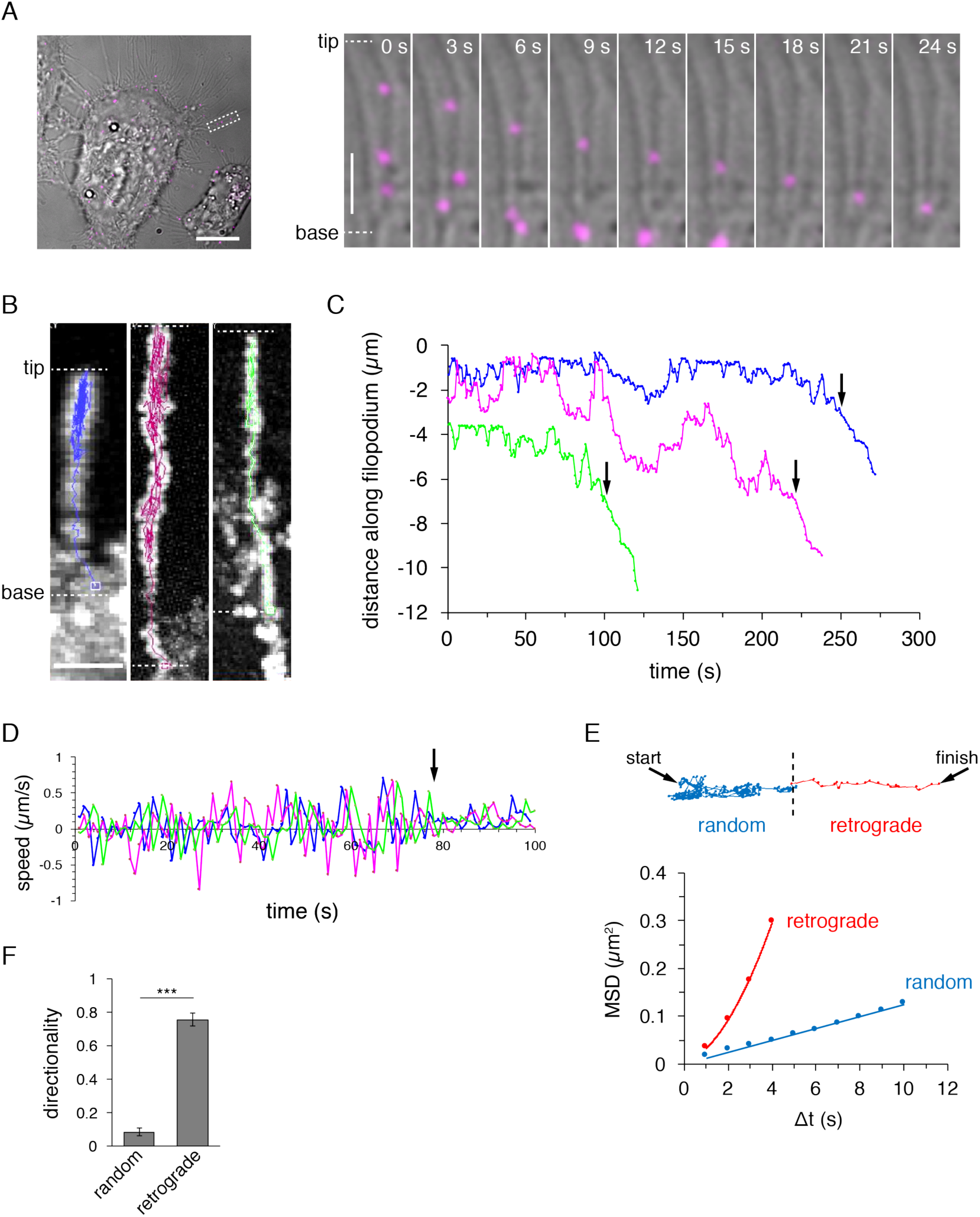
Ligand-bound IGF1R underwent directed centripetal movement on filopodia. Surface IGF1R on DU145 cells was labelled using biotin-IGF1 and streptavidin-QDs, before wide-field fluorescent imaging at 37°C. (A) *Left*: an overlay of IGF1-QDs (magenta) with the brightfield image (grayscale)—a representative image. Scale bar, 10 µm. *Right*: Centripetal transport of activated IGF1R on a filopodium at successive indicated times. Scale bar, 2 µm. (B) Directed transport on filopodia is preceded by non-directional movement. Three independent IGF1-QD trajectories (left) in blue, pink and green are shown overlaid onto the QD image stacks (grayscale). Scale bar, 2 µm. (C) Plot of distance along the filopodium against time for each of the IGF1-QDs in (B); arrows indicate where directed retrograde transport started. Scale bar, 2 µm. (D) Speed-time plots of the corresponding IGF1-QDs referring to the last 100s of imaging, before the IGF1-QD reached the filopodium base; speed is assigned a positive or negative sign depending if the QD is moving towards or away from the cell body, respectively. Arrow points to the onset of the retrograde movement of IGF1-QDs; tracks were aligned according to the onset of this movement transition. (E) In a plot of MSD against time, the non-directional part of the IGF1-QD trajectory (blue) was fitted linearly and the directional part (red) was fitted to a polynomial. The representative trajectory being analyzed is shown above, color-coded correspondingly, with its start and finish marked with arrows. (F) Directionality values differ significantly between the directed and random phase of IGF1-QD motion after ligand stimulation (mean ± s.e.m.; N = 8; *** P<0.001, unpaired t-test).

Here we report that IGF1 receptor gets cleared from filopodia after activation mediated by IGF1 binding, through an active transport toward the cell body. In contrast, on the apical plasma membrane of DU145 cells (data not shown) and MCF-7 cells (Koner et al., 2013), we have only observed Brownian diffusion of IGF1R, suggesting that this retrograde active transport is filopodium-specific. Our data point to myosin II-driven actin rearward flow as a probable mechanism for IGF1R trafficking on filopodia, in accordance with studies of viral particle transport and EGFR transport after ligand binding on filopodia (Lehmann et al., 2005; Lidke et al., 2004). In agreement, the average speed of ligand-bound IGF1R during the centripetal transport on filopodia corresponds to rearward actin flow rate in filopodia, reported in mouse neuroblastoma cells (Mallavarapu and Mitchison, 1999) How IGF1R could leverage rearward actin flow to move centripetally on filopodia remains to be established. We hypothesize that receptor activation triggers recruitment of unidentified receptor-binding effectors, which would mediate receptor tethering to F-actin in filopodia and consequently promote receptor transfer, as the receptor was not shown to bind actin directly. Directed transport of IGF1R on filopodia could be used to enable fine-tuned sensing of local IGF1 gradients and thereby regulate cell behavior spatially, by directing cell migration and tissue organization. The observed activation-dependent centripetal transport of IGF1R along the filopodia suggests a role in the adaptation of the sensing pathway by downregulating filopodial IGF1R in response to strong signals to help reset the system so that the cell is able to respond (again) to changes in ligand levels.

## Materials and Methods

### Reagents

Mouse (*Mus musculus*) monoclonal anti-IGF1R antibody (clone 24-31) recognizing the extracellular portion of IGF1R (Soos et al., 1992) was expressed and purified from the hybridoma, a kind gift of Ken Siddle (University of Cambridge). Affi_IGF1R_ (Li et al., 2010), the anti-IGF1R immunoglobulin-like binding protein bearing a C-terminal His_6_ tag and an N-terminal acceptor peptide (Beckett et al., 1999), was generated from pET21b-ZSPA (Holm et al., 2009) by inverse PCR with the forward primer 5’-AATCGAAAACAGTCTACCGCATTTATTTCTAGCCTTGAAGATGACCCAAGCCAAA GCGCT and the reverse primer 5’-TAGATTCGGTAATGCCAGGATTTCGATTGCAGCATAGAAACCTTCTTTGTTGAATTTGTTGTC. Affi_IGF1R_ was expressed in *E. coli* and purified as previously described for ZSPA (Holm et al., 2009). Affi_IGF1R_ was biotinylated site-specifically with BirA, before dialysis into PBS (Holm et al., 2009). Monovalent streptavidin (mSA) (Howarth et al., 2006) was expressed and purified as described (Howarth and Ting, 2008), before conjugating to Alexa Fluor 555 succinimidyl ester (Life Technologies, Carlsbad, US) following manufacturer’s instructions.

### Cell culture

DU145 human (Homo sapiens) prostate cancer cell-line (a kind gift of Val Macaulay, University of Oxford) and HeLa cells were grown in high-glucose DMEM with 10% fetal bovine serum (PAA, Pasching, Austria), 50 U/mL penicillin and 50 µg/mL streptomycin. Prior to experiments, cells were serum-deprived overnight by culturing in DMEM with 50 U/mL penicillin and 50 µg/mL streptomycin and then treated with inhibitors as follows: 120 nM AZ12253801 (Aleksic et al., 2010) (a kind gift from AstraZeneca, London, UK) for 1 h, 20 µM (-)-blebbistatin (Sigma-Aldrich, Gillingham, UK) for 1 h, 4 µM cytochalasin D (Sigma-Aldrich, Gillingham, UK) for 1 h, 80 µM dynasore (Sigma-Aldrich, Gillingham, UK) for 1 h, and 16 µM nocodazole (Sigma-Aldrich, Gillingham, UK) for 2 h. Cells were maintained in the medium with inhibitors until fixation. Live cell labeling was performed in PBS supplemented with 5mM MgCl_2_ (PBS-Mg) and 1% dialyzed BSA. pcDNA3 containing human IGF1R cDNA was a kind gift of Val Macaulay, University of Oxford. pECFP-H2B (human histone H2B fused to enhanced cyan fluorescent protein) has been described (Platani et al., 2002). pEGFP-N1 containing Lifeact-monomeric enhanced green fluorescent protein (Lifeact-mGFP) (Riedl et al., 2008) was a kind gift of Dan Davis, Imperial College. Transient transfections were performed with Turbofect (Fermentas, St. Leon-Rot, Germany) according to the manufacturer’s protocol.

### IGF1R time-course assays

DU145 cells growing on glass coverslips were serum-starved overnight and then either pre-treated with inhibitors or left untreated. The time-course was done by treating cells with 10 nM human des(1-3)IGF1 (ProSpec-Tany Technogene, Ness-Ziona, Israel) for 2, 5, 10 or 30 min at 37°C. Des(1-3)IGF1 binds like full-length IGF1 to IGF1R but avoids the complication of binding to IGFBPs (Ross et al., 1989). Control cells were either incubated with the inhibitor, an equal concentration of dimethyl sulfoxide (DMSO) to that received by the inhibitor-treated cells, or left untreated for the same time period. After incubating with des(1-3)IGF1, cells were fixed with 3% (w/v) paraformaldehyde (Sigma-Aldrich, Gillingham, UK) for 20 min at 37°C.

### Immunofluorescence

Following fixation, paraformaldehyde was quenched with 50 mM ammonium chloride in PBS for 10 min at room temperature, and samples were blocked with 3% BSA in PBS for 30 min. Surface IGF1R was labelled with 24-31 antibody at 4 µg/ml, followed by 10 µg/ml AlexaFluor 647-conjugated goat anti-mouse antibody (Life Technologies, Carlsbad, US), both for 1 h at room temperature. Cell membrane labeling was done for 10 min using 5 µg/ml WGA-AlexaFluor 488 (Life Technologies, Carlsbad, US) in PBS. Samples were mounted onto microscope slides using Prolong Gold Antifade reagent (Life Technologies, Carlsbad, US).

### IGF1R labelling in living cells and time-lapse imaging

IGF1R labelling in living cells was done as described (Koner et al., 2013), with some modifications. Briefly, DU145 cells growing in glass-bottom 35-mm imaging dishes (Willco, Amsterdam, Netherlands) were serum-starved overnight, rinsed with PBS-Mg and treated with 5 nM biotinylated human des(1–3) IGF1 (IBT, Binzwangen, Germany) or 5 nM biotin-Affi_IGF1R_ in PBS-Mg containing 1% dialyzed BSA for 5 min. Des(1–3) IGF1 is a modified IGF1 with normal binding to IGF1R and minimized interaction with IGF binding proteins (IGFBPs) (Ross et al., 1989). After three washes in PBS-Mg, 1 nM Qdot605-streptavidin (Life Technologies, Carlsbad, US) in PBS-Mg containing 1% dialyzed BSA was added to cells and incubated for 5 min. After three washes with PBS-Mg, L-15 Leibovitz medium (Life Technologies, Carlsbad, US) pre-warmed to 37°C was added and imaging started immediately. Time-lapse movies (20 min duration; 1 s between frames) were taken using an incubated (37°C) wide-field DeltaVision Elite fluorescence microscope (Applied Precision, Washington, US) with 100× objective and optovar lens (1.6×), using softWoRx 5.0.0 software (Applied Precision, Washington, US). QDs were imaged with 490DF20 excitation, 617DF73 emission, and a Chroma 84100bs polychroic filter set using 0.2 s exposure. AlexaFluor 647 was imaged with 640DF20 excitation, 685DF40 emission, and a Chroma 84100bs polychroic filter set with 2 s exposure. AlexaFluor 488 was imaged with 490DF20 excitation, 528DF38 emission, and a Chroma 84100bs polychroic filter set with 0.1 s exposure. AlexaFluor 568 was imaged with 555DF28 excitation, 617DF73 emission, and a Chroma 84100bs polychroic filter set with 2 s exposure.

### Affi_IGF1R_ binding specificity

HeLa cells were transiently co-transfected with pcDNA3-IGF1R and Lifeact-mGFP plasmids and serum-starved overnight. The transfected cells were incubated with 40 µM biotin-Affi_IGF1R_ in PBS-Mg with 1% dialyzed BSA for 30 min on ice. After three washes with PBS-Mg, cells were treated with 200 nM mSA-AlexaFluor 555 in PBS-Mg with 1% dialyzed BSA for 15 min on ice, rinsed as above and imaged immediately. To assess biotin-IGF1 binding specificity, HeLa cells were transiently co-transfected with pcDNA3-IGF1R and H2B-CFP plasmids, or untransfected. After serum-starving overnight, cells were incubated with 50 nM biotin-IGF1 in PBS-Mg with 1% dialyzed BSA for 10 min on ice. After three washes with cold PBS-Mg, 200 nM mSA-AlexaFluor 555 in PBS-Mg with 1% dialyzed BSA was added for 10 min on ice. Cells were washed with cold PBS-Mg and imaged as soon as possible. To further validate the specific binding of biotin-IGF1 to IGF1R, live DU145 cells were either pre-incubated with 55 µM non-biotinylated Affi_IGF1R_ in PBS-Mg with 1% dialyzed BSA for 15 min at room temperature to block IGF1-binding on IGF1R, or left untreated. Then, cells were treated with 5 nM biotin-IGF1 in PBS-Mg with 1% dialyzed BSA for 5 min at room temperature, rinsed three times with PBS-Mg, and incubated with 1 nM Qdot605-streptavidin in PBS-Mg with 1% dialyzed BSA for 5 min at room temperature. The cells were placed on ice and imaged as soon as possible.

### Image analysis and processing

All images were processed in ImageJ (v.1.46r; National Institutes of Health, USA). Analysis was done on fully spread, non-mitotic DU145 cells. Filopodia were identified based on their appearance in the WGA or differential interference contrast (DIC) channel, as unbranched, thin processes of uniform width (≤0.3 µm) along the axis, emanating from the cell periphery. The length of filopodia on DU145 cells varied from 1 to 12 µm while most of the values were in the 2-6 µm range; processes shorter than 1 µm were not analyzed. Time-lapse movie analysis was performed on stable filopodia of constant length throughout the movie. For time-course assays, RGB images were color-separated, converted to 8-bit grayscale images and background subtracted (ImageJ). Gaussian blur (radius 1; ImageJ) was applied in the representative images. The regions of interest (ROI) (filopodia) were selected using a 5-pixel (0.33 µm) wide segmented line, which was carefully overlaid onto filopodia in the WGA channel. ROI was quantified in the AlexaFluor 647 channel using ImageJ routines for measuring length, mean fluorescence intensity, and area of the selection. Filopodial IGF1R signal was calculated as the quotient of the mean IGF1R fluorescence intensity and the area of the selection. Data are expressed as the mean and the standard error of the mean (s.e.m.). Statistical analysis was done in GraphPad Prism v. 5.02 (GraphPad Software, San Diego, US) and consisted of one-way analysis of variance (ANOVA) followed by post-tests to find which means are significantly different from one another (Tukey’s test was used to compare every pair of means, and Dunnett’s test was used to compare every mean to a control mean). A p-value of 0.05 or less was considered statistically significant. Time-lapse movies were converted from 16-bit to 8-bit grayscale image stacks and processed using the fast Fourier transform (FFT) bandpass filter in ImageJ, to improve the signal-to-noise ratio. The processed image stacks were analyzed as described (Koner et al., 2013). Briefly, View5D, an ImageJ plug-in developed by R. Heintzmann (King’s College, London), was used for semi-automatic tracking of calculated centers of intensity in each frame. The quality of the tracking was verified visually for each frame. Coordinates, Total Speed, Average Speed, Directionality index and mean square displacement (MSD) values were obtained from the View5D output file. The Total Speed was calculated as the distance between the first and the last point of measurement divided by the total measurement time. The Average Speed was calculated as the mean of the speed of movement in each frame. Directionality Index (the Total Speed divided by the Average Speed) is invariant under rotation and ranges from 0 to 1, where index of 1 signifies completely directed movement (along a straight line and only forward), whereas an index of zero means that the particle returned to the origin. Diffusion coefficients (D) were acquired by either fitting the whole linear MSD curve (normal diffusion) or the linear region (first 4-10 points) of the MSD curves (for anomalous and confined diffusion) with the equation *MSD=2D*Δ*t* (for one-dimensional filopodial geometry). The unpaired t-test was done in GraphPad Prism software.

## Acknowledgements

We thank Val Macaulay for constructs, and for sharing the IGF1R inhibitor. We thank Danijela Vignjevic, Evelyne Coudrier, Tamara Aleksic and Rafael Galupa for critical reading of the manuscript.

## Author Contributions

D.K conceived the study and designed all experiments. Experiments were performed by D.K and M.F. All recombinant proteins were produced by M.F. Data analysis was done by D.K. D.K wrote the manuscript.

