## Supplementary Materials for "IGF1R undergoes active and directed centripetal transport on filopodia upon receptor activation"

**This PDF file includes:**

Supplementary figures S1 and S2

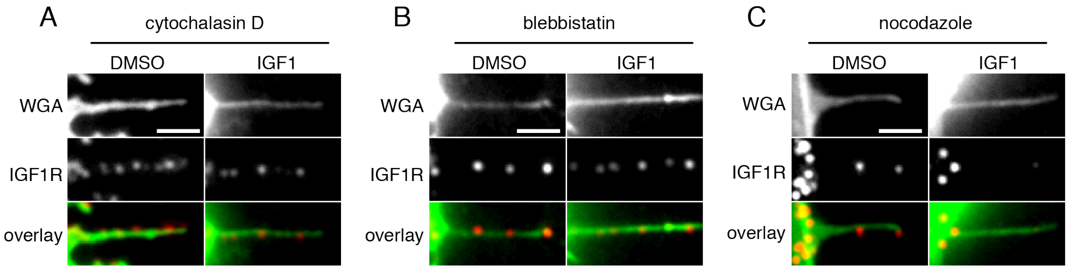


**Fig. S1. IGF1-induced clearance of IGF1R from filopodia is dependent on actin polymerization and myosin II activity, but does not require microtubule stability.** DU145 cells were serum-starved overnight and then treated with cytochalasin D (A), blebbistatin (B), nocodazole (C), or DMSO (vehicle control), before adding IGF1 for 30 min and staining for IGF1R intensity on filopodia. Representative images are showing filopodia stained with wheat germ agglutinin (WGA) (top) and anti-IGF1R antibody (middle), with the merge shown on the bottom (WGA in green, IGF1R in red). Scale bar, 5 µm.

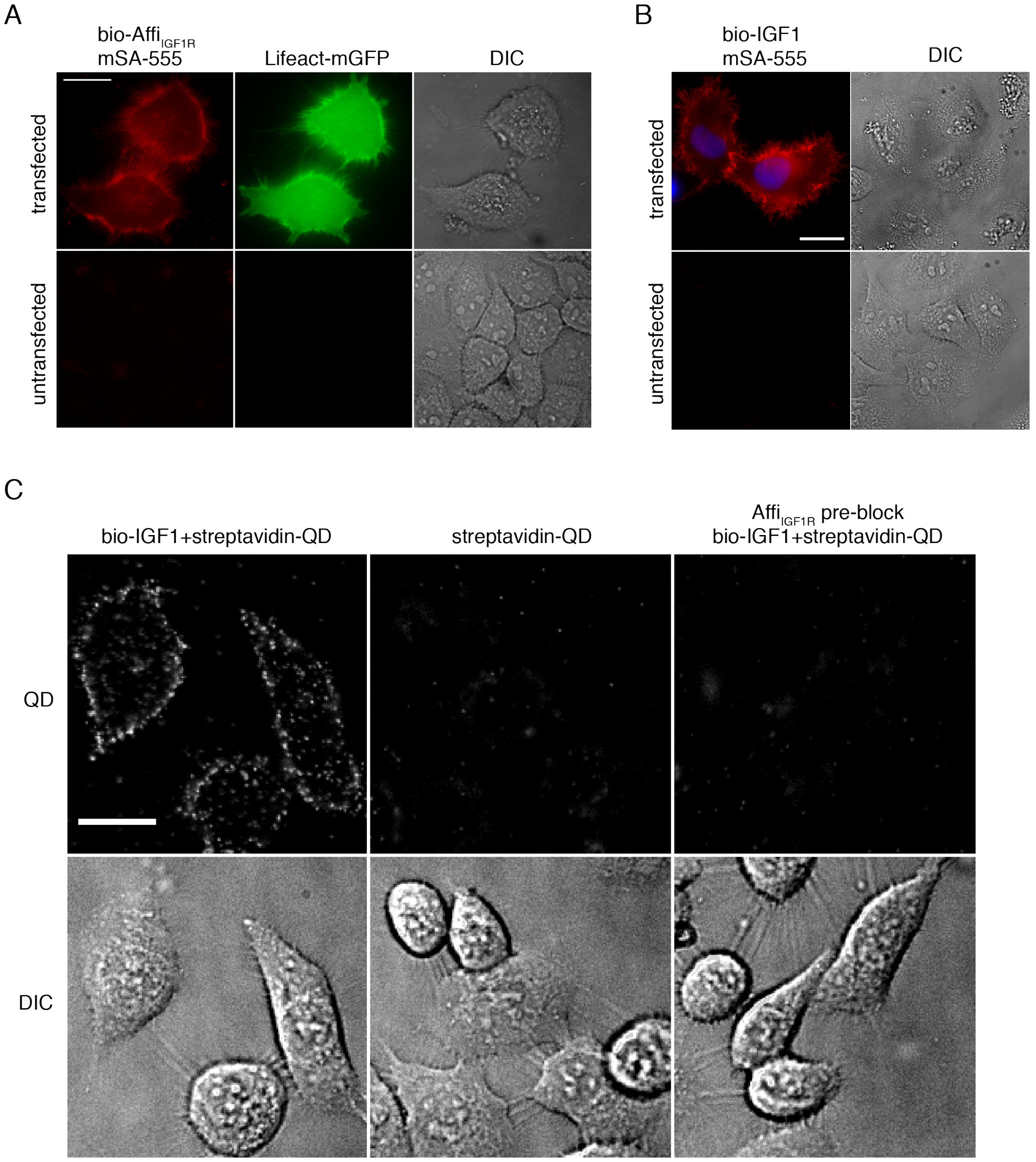


**Fig. S2. Specificity of binding of Affi$_{IGF1R}$ and bio-IGF1.** (A) Affi$_{IGF1R}$ binding specificity. HeLa cells were transfected with IGF1R, and Lifeact-mGFP as a cotransfection marker (top), or untransfected (bottom). After serum-starving overnight, the cells were labeled with biotinylated Affi$_{IGF1R,}$ followed by monovalent streptavidin-AlexaFluor 555 (mSA-555) and imaged on an epifluorescence microscope. mSA-555 (red), Lifeact-mGFP (green) and DIC images (grayscale) images are shown. Scale bar, 20 μm. (B) Bio-IGF1 binding to IGF1R validation. HeLa cells were transfected with IGF1R, using H2B-CFP as a co-transfection marker (top), or left untransfected (bottom). After serum-starving overnight, the cells were labelled with bio-IGF1 followed by mSA-555. Merged mSA-555 (red) and H2B-CFP (blue) images are shown, as well as DIC images (greyscale). Scale bar, 20 μm. (C) Bio-IGF1 binds specifically to IGF1R. DU145 cells were labelled with bio-IGF1 followed by streptavidin-QD (left). In the middle panel, bio-IGF1 was omitted. In the right panel, cells were pre-treated with

Affi$_{IGF1R}$ to establish whether any of the bio-IGF1 binding sites were distinct from IGF1R. QD (top) and DIC (bottom) images from wide-field microscopy are shown, both in grayscale. Scale bar, 20 μm.
